# Nonessential nucleoporins Nup2, Nup60 and Nup84 are required for normal meiosis in *Saccharomyces cerevisiae*

**DOI:** 10.1101/054791

**Authors:** Daniel B. Chu, Sean M. Burgess

## Abstract

The nuclear pore complex (NPC) selectively transports cargo between the nucleus and the cytoplasm. The inner nuclear membrane (INM) face of the NPC also serves as a hub where gene silencing and DNA repair are spatially coordinated. In *Saccharomyces cerevisiae*, partitioning of active and silenced chromatin at subtelomeric regions depends on the boundary activity of nonessential nucleoporin proteins Nup2 and Nup60, along with Htz1, the histone variant H2A.Z. The INM is also important for the chromosome events of meiosis since Ndj1-mediated telomere attachment and clustering at the INM is required for efficient homolog pairing, recombination, and segregation. Here we tested possible meiotic roles for Nup2, Nup60, Htz1, and other nonessential nucleoporins by analyzing the effects of deletion mutations on sporulation, spore viability and possible phenotypic epistasis with *ndj1Δ*. Deleting *NUP2*, *NUP60*, and *HTZ1* reduced spore viability compared to wild-type (WT). A detailed analysis of spore lethality indicated that homolog nondisjunction in these mutants was elevated compared to WT, yet unlike *ndj1Δ*, this was not the predominant cause of spore death. Deleting *NUP84* reduced meiosis I nuclear divisions, while deleting *NUP53*, *NUP100*, and *NUP157* had no effect on sporulation, spore viability, or the kinetics of meiosis I progression. Surprisingly, *nup2Δ ndj1Δ* uniquely failed to undergo meiosis I nuclear divisions, suggesting Nup2 and Ndj1 function in partially redundant pathways or create a poisonous intermediate. The meiosis I division was also delayed by 2 hours in *nup2Δ* compared to WT pointing to a specialized role for Nup2 in the meiotic program.

## Introduction

The nuclear pore complex (NPC) is a ring structure, which sits in the nuclear envelope bridging the inner and outer nuclear membranes. The primary function of the NPC is to selectively transport cargo into and out of the nucleus. Aside from its nuclear transport role the NPC has also been implicated in aspects of nuclear architecture that impact processes such as the DNA damage response, transcriptional silencing, boundary activity, transcriptional memory, mRNA processing, and mRNA export (Fahrenkrog *et al.* 2000; Denning *etal.* 2001; Dilworth *et al.* 2005; Bukata *et al.* 2013; Ptak *et al.* 2014). The NPC is composed of proteins called nucleoporins (Nups), which fall under several functional components of the NPC. The main structure the NPC is made up of two inner rings which are sandwiched between two outer rings, with one outer ring touching the nucleoplasmic side and the other touching the cytoplasmic side (Figure 1; Aitchison and Rout 2012).

**Figure 1.**
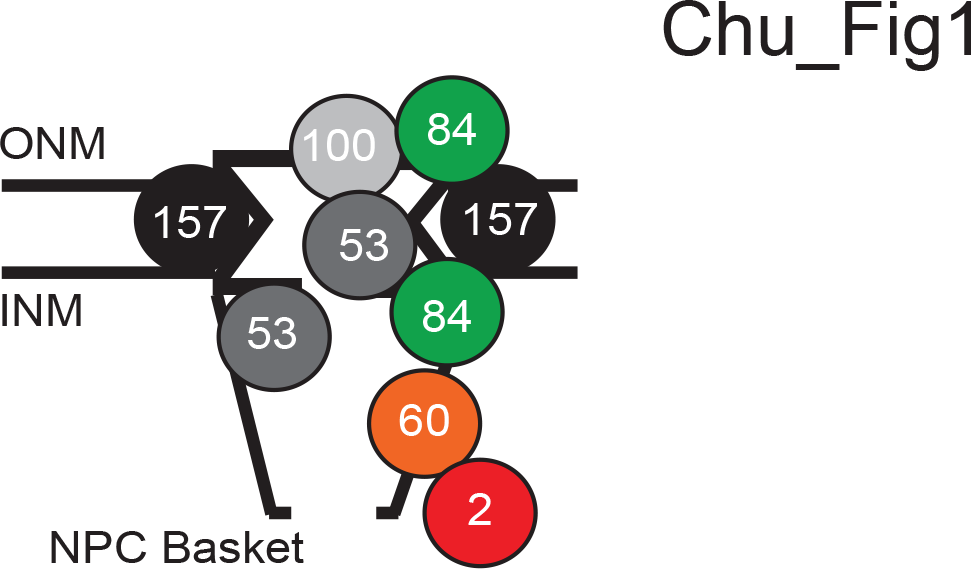
**Schematic of the position(s) of each nonessential Nup protein examined in this study** Z-axis slice schematic of the nuclear pore complex showing the relative position of the Nups analyzed in this study (adapted from Alber et al., 2007). The cytoplasmic side is on the top of the outer of the nuclear membrane (ONM) and the nucleoplasmic side in below the inner nuclear membrane (INM; thick horizontal lines). Nup84 is located in the outer ring (green); Nup157 is located in the inner ring (black); Nup100 is an FG-Nup found on the cytoplasmic side of the pore channel (light gray); Nup53 is an FG-Nup found on both sides of the channel (dark gray); Nup60 (orange) and Nup2 (red) are FG-Nups is primarily on the nucleoplasmic side of the channel near the nuclear basket.

The selective permeability of the NPC is generated by phenylalanine-glycine (FG) Nups which contain an FG-repeat motif made up of FGs separated by ~20 amino acids (Aitchison and Rout 2012). FG Nups predominantly attach to the inner rings and the FG-repeat motif generally results in a long unstructured filament which extends into the central channel of the NPC (Alber *et al.* 2007). FG Nups filaments also extend into the cytoplasm as cytoplasmic filaments and into the nucleoplasm as the nuclear basket. FG Nups serve to block passive diffusion and direct transport across the NPC (Figure 1). The cytoplasmic filaments and nuclear basket serve to direct release of cargo following transport through the NPC.

Indirect evidence from non-yeast species links Nups to the meiotic program which is a specialized series of cell divisions required to create haploid gametes from diploid parent cells (Zickler and Kleckner 1998; Zickler and Kleckner 2015). A hallmark of meiosis is the pairing, synapsis and recombination between homologous chromosomes that ensures their segregation at the first meiotic division (Zickler and Kleckner 2015). A mutation in Tmem48/NDC1 in mouse causes defects in homologous chromosome pairing and the arrest of cells in late meiotic prophase, and sterility (Akiyama *et al.* 2013). In *Drosophila melanogaster,* a P element insertion into Nup154 (yeast *NUP157*) results in defects in both male and female meiosis with complete prophase arrest in males (Gigliotti *et al.* 1998). The mouse Nup2 homolog, mNup50, and its paralog, n50rel, are enriched in the testis (Fan *et al.* 1997; Smitherman *et al.* 2000).

During meiotic prophase, telomeres cluster at the nuclear envelope near the microtubule organizing center in a characteristic bouquet conformation (Zickler and Kleckner 1998). In plants and animals, nuclear pores reorganize to cluster near the bouquet telomeres during meiotic prophase; in yeast, direct interactions between bouquet telomeres and nuclear pores have been observed (Church 1976; Holm 1977; Klein *et al.* 1992). In yeast, the bouquet is directed by Ndj1 which physically connects the telomeres to the nuclear envelope (Chua and Roeder 1997; Conrad *et al.* 1997).

In this study, we explored possible role(s) for nonessential Nups in mediating normal meiotic progression in budding yeast. Through a past genome duplication event in budding yeast, deletions of several *NUP* genes are viable due, in part, to functional redundancy (Kellis *et al.* 2004; Aitchison and Rout 2012). We created deletions in genes encoding a component of the inner ring (Nup157), the outer ring (Nup84), and four FG Nups (Nup2, Nup53, Nup60, and Nup100) that localize within the central channel or near the nuclear basket of the NPC (Wente *et al.* 1992; LOEB *et al.* 1993; AITCHISON *et al.* 1995; SINIOSSOGLOU *et al.* 1996; MARELLI *et al.* 1998; Rout *et al.* 2000). For this study, each deletion strain was tested for growth under different challenged regimens, sporulation efficiency, spore viability, and the timing of meiotic progression. We also evaluated possible genetic interactions between these mutations and the deletion of *NDJ1*.

## Results

### Growth of *nup* mutants in the SK1 yeast strain background

To evaluate the effects of *nup* mutations in meiosis, we created homozygous gene knockouts of the nonessential Nups listed above in the SK1 strain background, which is well known for its high efficiency of sporulation and spore viability (Figure 1; Kane and Roth 1974; Elrod *et al.* 2009). Since growth defects could contribute to inefficient meiosis, we analyzed the colony formation of each mutant and wild type (WT) strain at various temperatures on rich (YPD) and minimal solid media. At 20°, 25°, and 30°, all strains gave similarly sized colonies as WT except *nup60Δ* and *nup84Δ*, which displayed modest growth defects at these temperatures on YPD (Figure 2A). Surprisingly, *nup2Δ* and *nup60Δ* mutants gave larger colonies on YPD at 37° compared to WT (Figure 2A). The *nup100Δ* mutant also gave larger colonies compared to WT on minimal medium at all temperatures (Figure 2B). These results uncover surprising antagonistic pleiotropic phenotypes of *nup2Δ*, *nup60Δ* and *nup100Δ* mutants on colony growth.

**Figure 2.**
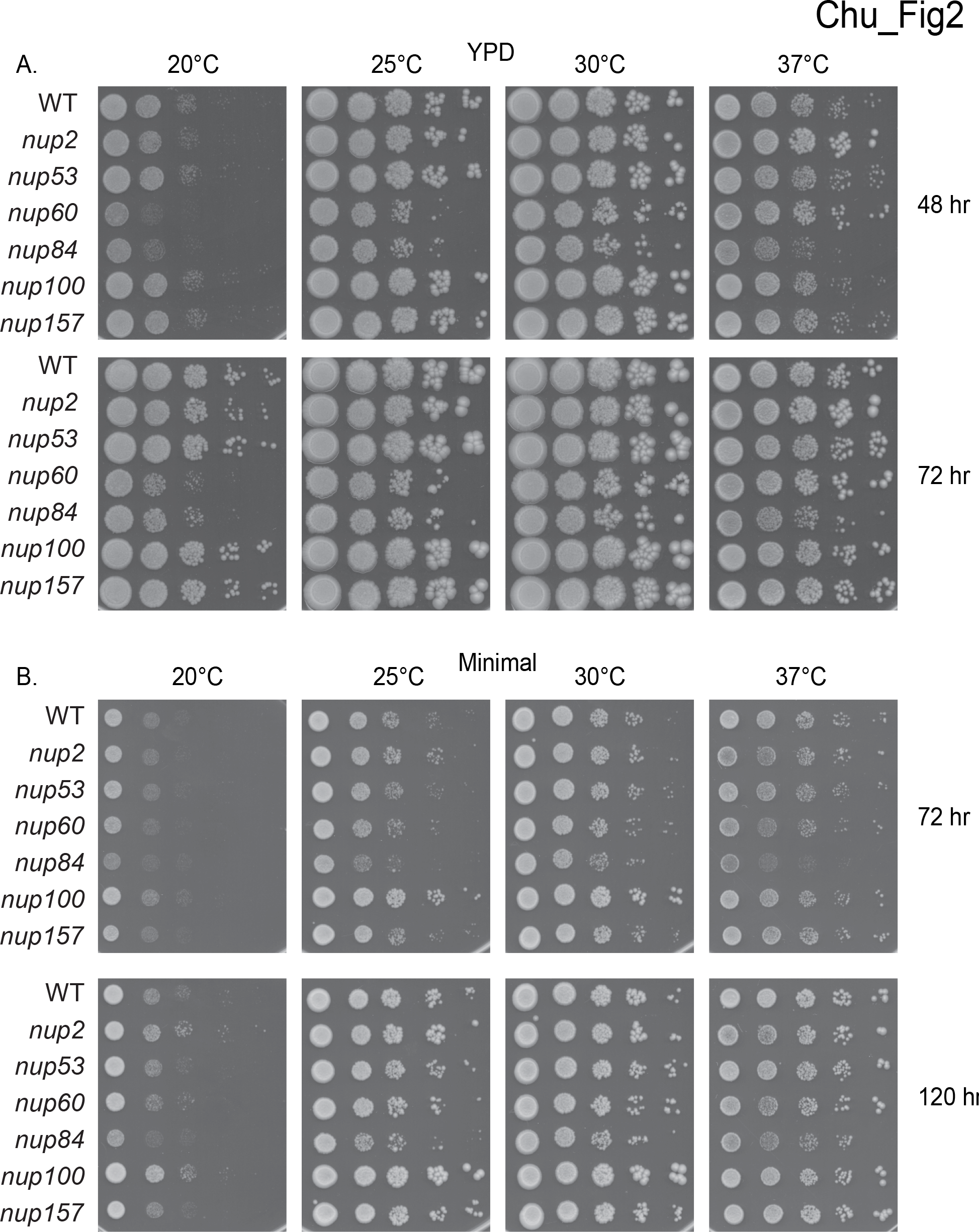
**Growth of nup mutants on rich media (YPD) and minimal media at 20°, 25°, 30°, and 37°**.(A) Growth of WT and *nup* mutants at on YPD after 48 and 72 hours at the indicated temperatures. Shown are spotted 1:10 serial dilutions. (B) Same as in (A) except cells were spotted on minimal medium and imaged after 72 and 120 hours at the indicated temperatures.

### Sporulation efficiency and spore viability of *nup* mutants

We next assessed sporulation and spore viability in the *nup* mutants at%30° (Table 1). All *nup* mutants sporulated with the same efficiency as WT (95%; n = 200) on solid sporulation medium except for *nup60Δ* (71%; n = 200) and *nup84Δ* (<1%; n = 200). While 70% of *nup84Δ* cells could undergo the meiosis I (MI) nuclear division, no cells went on to form tetrads. For the remaining strains, four-spore tetrads were dissected on YPD and growth was assessed after four days of incubation at 30°. Among four-spore tetrads, WT strain gave 97.1% (n = 1152) viable spores from four-spore asci, while *nup2Δ* and *nup60Δ* gave 86.1% (n = 1536) and 70.5% (n = 1056), respectively (Table 1). There was no measurable defect in spore viability in *nup53Δ* (95.8%; n = 288), *nup100Δ* (98.6%, n = 288) or *nup157Δ* (98.3%; n = 288; Table 1).

**Table 1.** Sporulation and spore viability in *nup* mutants

| Strain | % Sporulation efficiency (n) | Tetrads dissected | 4-sv | 3-sv | 2-sv | 1-sv | 0-sv | % Spore viability |
| --- | --- | --- | --- | --- | --- | --- | --- | --- |
| WT | 95% (200) | 288 | 269 | 10 | 6 | 1 | 2 | 97.1 |
| WT<sup>a</sup> |  | 1199 | 1087 | 77 | 32 | 1 | 2 | 96.8 |
| *nup2Δ* | 92% (200) | 384 | 239 | 95 | 36 | 10 | 4 | 86.1 |
| *nup60Δ* | 71% (200) | 264 | 85 | 96 | 46 | 25 | 12 | 70.6 |
| *nup84Δ* | <1% (200) | NA |  |  |  |  |  |  |
| *nup53Δ* | 94% (200) | 72 | 66 | 3 | 1 | 1 | 1 | 95.8 |
| *nup100Δ* | 93% (200) | 72 | 69 | 2 | 1 | 0 | 0 | 95.9 |
| *nup157Δ* | 94% (200) | 72 | 69 | 1 | 2 | 0 | 0 | 98.2 |
| *htz1Δ* | 67% (200) | 288 | 77 | 91 | 69 | 40 | 11 | 66.0 |
| *ndj1Δ* | 75% (200) | 192 | 136 | 14 | 19 | 6 | 17 | 82.0 |
| *ndj1Δ*<sup>a</sup> |  | 851 | 482 | 112 | 123 | 38 | 96 | 74.9 |
| *nup2Δ ndj1Δ* | 1% (200) | NA |  |  |  |  |  |  |
| *nup60Δ ndj1Δ* | 46% (200) | 48 | 16 | 4 | 8 | 3 | 7 | 65.1 |
| *nup53Δ ndj1Δ* | 75% (200) | 144 | 103 | 10 | 12 | 6 | 13 | 82.0 |
| *nup100Δ ndj1Δ* | 74% (200) | 72 | 48 | 3 | 14 | 1 | 6 | 79.9 |
| *nup157Δ ndj1Δ* | 72% (200) | 72 | 39 | 5 | 14 | 4 | 10 | 70.5 |
| *htz1Δ ndj1Δ* | 41% (200) | 72 | 15 | 18 | 17 | 13 | 9 | 55.9 |

<sup>a</sup> Data reported in Wanat et al., 2008

We included the deletion of *HTZ1* (histone variant H2A.Z), since Htz1 is associated with the NPC and plays a role in the boundary activity of Nup2 and Nup60 (Dilworth *et al.* 2005). The *htz1Δ* mutation gave 65.9% viable spores among four-spore tetrads (n = 1152; Table 1). Since Nup2, Nup60, and Htz1 function in telomere silencing, we tested their mutant phenotypes in the *ndj1Δ* mutant strain background, in which telomere attachment to the nuclear envelope is disrupted (Trelles-Sticken *et al.* 1999). The *ndj1Δ* mutant alone gives 82.0% (n = 768) spore viability in our strain background compared to 75% as reported by Wanat et al., 2008. The *nup60Δ ndjlΔ* and the *htzlΔ ndjlΔ* double mutants showed no epistasis with respect to spore inviability compared to the single mutant phenotypes, suggesting that Nup60 and Htz1's roles during meiosis are independent of Ndj1 (Table 1). Interestingly, the *nup2Δ ndj1Δ* double mutant failed to undergo the MI nuclear division. A more detailed analysis of this phenotype will be reported separately (D.B.C and S.M.B, unpublished).

### MI progression is delayed in *nup2Δ* compared to WT

We measured the timing of the MI nuclear division in the *nup* knockout strains following the transfer of synchronized cell cultures to sporulation medium (t = 0 hours). Post-MI stages are marked by the presence of at least two DAPI-stained nuclear bodies. In *nup2Δ,* MI was delayed ~two hours compared to WT (Figure 3). By contrast, *nup53Δ*, *nupl00Δ*, and *nupl57Δ* deletion mutations had no measurable effect on MI timing. The *nup60Δ*, *nup84Δ*, and *htz1Δ* mutants were excluded from the MI analysis because the cells would be unlikely to enter meiosis synchronously in liquid culture due to their mitotic growth defects at 30°.

**Figure 3.**
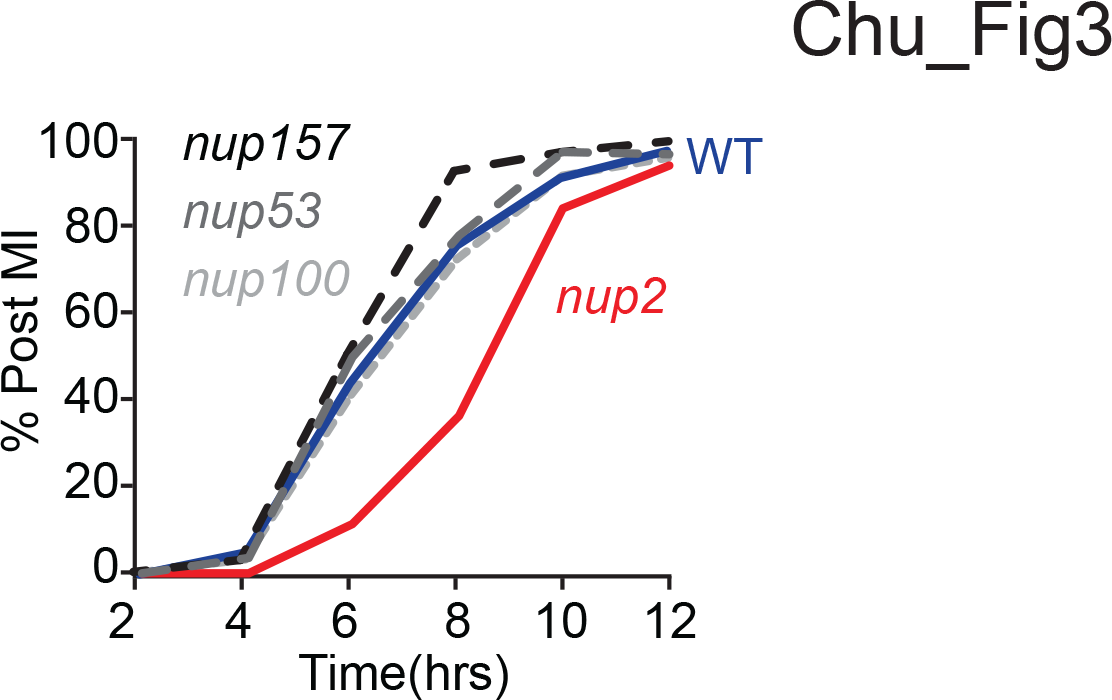
**Meiotic phenotypes of nonessential nucleoporins** (A) Percent viable spores are based on growth of spore clones from dissected 4-spore tetrads for the indicated deletion mutants (n = number of total spores dissected) after four days at 37C. (B) Representative meiotic time course of WT (blue), *nup2Δ* (red), *nup53Δ* (dark gray), *nup100Δ* (light grey), and *nup157Δ* (black). Multinucleate cells are those with 2 or more DAPI staining bodies. At least 200 cells were scored for each data point. (C) Same as in part (A) for the indicated deletion strains.

### Causes of spore death in *nup2Δ*, *nup60Δ*, and *htzlΔ*

The distribution of tetrads giving 4, 3, 2, 1, or 0-viable spores can lend insight into the causes of spore death (Chu and Burgess 2016). For example, mutants with demonstrated increased levels of MI nondisjunction (MI-ND) also give rise to an increased incidence of two spore and zero spore viable tetrads (Shonn *et al.* 2000; Martini *et al.* 2006; Wanat *et al.* 2008). We recently introduced two R-based tools, TetSim and TetFit, for analyzing the causes of mutant spore death based on the distribution of live:dead tetrad classes (File S1; Chu and Burgess 2016). Detailed annotated R-scripts and examples of all the generated outputs for each test are presented in File S1.

TetSim simulates the expected distribution of tetrads if spore death was purely random; that is, when one spore dies there is no corresponding death of another spore (Chu and Burgess 2016). If spore inviability of a given mutant was due solely to random spore death (RSD), the observed distribution of live:dead tetrads should resemble the output of TetSim (Figure 4A). If spore death was due, in part, to MI-nondisjunction, the observed distribution would show outliers from the 99% confidence limits of the simulated distribution. Applying TetSim to previously published tetrad data sets for WT and *ndj1Δ* (Wanat et al., 2008) using TetSim showed that not all of the live:dead tetrad classes fit a distribution predicted by RSD alone (Figure 4A, horizontal solid lines; Chu and Burgess 2016). Our own tetrad data agreed with these findings. For *nup2Δ, nup60Δ,* and *htz1Δ*, at least two of the live:dead tetrad classes fell outside the 99^th^ percentile for RSD alone (Figure 4A). These results indicate that the increased levels of spore death in these mutants was not solely due to random spore death. The *nup53Δ, nup100Δ*, and *nup157Δ* mutants were not analyzed since spore viability in these strains was indistinguishable from WT (Table 1).

**Figure 4.**
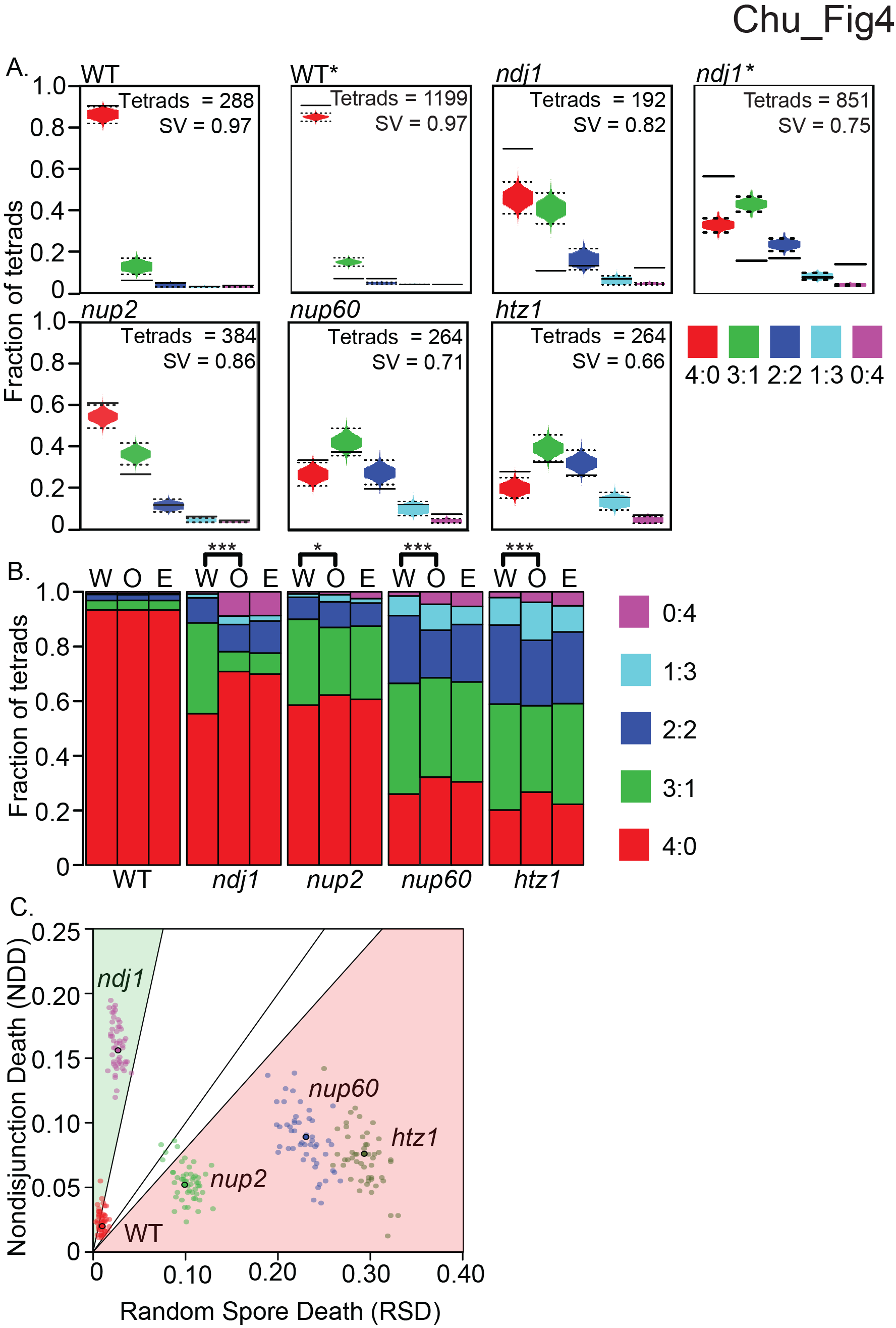
Estimation of nondisjunction death and random spore death caused by *nup2Δ*, *nup60Δ*, and *htz1Δ*. (A) TetSim simulates a distribution of 4:0, 3:1, 2:2, 1:3, and 0:4 tetrads based on the observed fraction of viable cells (Chu and Burgess 2016) and compares the 99% cutoff values for the simulated distribution to the observed distribution for each genotype. The number of simulations for each panel is equal to the number of tetrads dissected experimentally. A distribution of dissected tetrad simulations is shown (in red, green, blue, teal, and purple) with the upper and lower dashed lines represent 99% and 1% percentiles, respectively. The observed frequencies of live:dead tetrad distributions for individual strains are shown as solid lines. Comparison of simulated tetrad distributions with mutant data from previously published data sets for WT and *ndj1Δ* (Wanat *et al.* 2008). Note the fraction of viable spore in the *ndj1Δ* mutant was higher than previously reported by Wanat et a., 2008. (B) Best fitting distributions of 4 (red), 3 (green), 2 (blue), 1 (teal), and 0 (purple)-viable spore tetrads in *nup2Δ*, *nup60Δ*, *htz1Δ*, and *ndjlΔ* mutants generated by TetFit (Chu and Burgess, 2016). The observed tetrad distributions of genotype (O, middle bar), the expected tetrad distributions with the best fitting MI-ND and RSD (E, right bar), and the expected tetrad distributions using the WT MI-nondisjunction rate and the best fitting RSD (W, left bar). Significance is noted as * *P* < 0.05; *** *P* < 0.001 (chi-squared, Holm corrected). (B) Output of TetFit, which finds the calculated estimated distributions of each live:dead tetrad class. The observed tetrad distributions of genotype (O, middle bar), the expected tetrad distributions with the best fitting MI-ND and RSD (E, right bar), and the expected tetrad distributions using WT MI-ND and the best fitting RSD (W, left bar). (C) 50 simulations of TetFit using dissected tetrad simulations corresponding to the estimated tetrad distributions shown in part (B) and the number of experimental tetrads shown in part (A). The solid line represents a NDD/RSD slope of 1, which indicates an equal contribution of NDD and RSD on spore inviability. The green shaded area covers the Class A mutants that have an NDD/RSD ratio above 3.3, and the pink shaded area covers Class B mutants that have an NDD/RSD ratio below 0.8 (Chu and Burgess 2016).

We next used the R-script TetFit to test if the decrease in observed spore viability for each genotype (Table 1 and Table S1) could be explained by MI-ND or a combination of MI-ND and RSD (Table S2; Chu and Burgess 2016). TetFit-A finds the combination of these values that gives the best fit to the observed tetrad distribution (Table 2; Figure 4B). For WT, *ndj1Δ*, *nup2Δ*, *nup60Δ*, and *htz1Δ* the calculated best fits were not statistically different from the observed distributions (*P* > 0.05, adjusted for multiple comparisons), suggesting that both MI-ND and RSD could account for all of the observed spore death (Figure 4B).

**Table 2.** Mean and standard deviation of 50 simulations of TetFit. Test

| Strain | RSD<sup>a</sup> | NDD<sup>a</sup> |
| --- | --- | --- |
| WT | 0.010 ± 0.003 | 0.024 ± 0.009 |
| *ndj1Δ* | 0.0262 ± 0.005 | 0.160 ± 0.018 |
| *nup2Δ* | 0.104 ± 0.011 | 0.054 ± 0.011 |
| *nup60Δ* | 0.229 ± 0.019 | 0.087 ± 0.023 |
| *htz1Δ* | 0.292 ± 0.017 | 0.072 ± 0.025 |

<sup>a</sup> Mean and standard deviation of estimated fraction of spores that die due to RSD (random spore death) and NDD (death due to nondisjunction) for each genotype.

TetFit also reports on the frequencies of death due to MI nondisjunction (nondisjunction death; NDD), thus allowing for comparison of the relative contributions of MI-ND and RSD to spore inviability. The NDD/RSD ratios for our WT and *ndj1Δ* data sets were 2.0%/0.9% and 15.78%/2.7%, respectively (Table 2 and Figure 4B); in both cases the cause of spore death was skewed to NDD. This is consistent with NDD/RSD outcomes of TetFit analyisis of previously published data sets for these genotypes (Wanat et al. 2008; Chu and Burgess, 2016), where the calculated NDD/RSD ratios for WT and *ndj1Δ* were 1.8%/1.8% and 20.8%/5.8%, respectively. The output of TetFit applied to the observed tetrad distributions of *nup2Δ*, *nup60Δ*, and *htz1Δ* gave greater NDD compared to WT, suggesting that spore death in these mutants can be attributed, in part, to MI-ND. However, the best fit NDD/RSD ratios in these mutants were skewed to greater RSD values: *nup2Δ* was 5.2%/9.9%; *nup60Δ* was 8.9%/23.1%; and *htzlΔ* was 7.5%/29.4% (Table 2 and Figure 4B). Thus, unlike *ndj1Δ*, the cause(s) of spore death in *nup2Δ*, *nup60Δ*, and *htz1Δ* appear to be predominantly due to RSD rather than MI-ND.

The magnitude of the NDD/RSD ratio defines two classes of mutants (Chu and Burgess, 2016. Class A mutants give NDD/RSD > 3.3 and include those with known defects in proper homolog segregation (e.g. *mad1Δ*, *mad2 Δ*, hypomorphic alleles of *spo11*, *ndj1Δ*, *csm4Δ*, *msh5Δ*, *mlh1Δ* and *exo1Δ* (Chua and Roeder 1997; Conrad *et al.* 1997; Kirkpatrick *et al.* 2000; Shonn *et al.* 2000; Marston *et al.* 2004; Cheslock *et al.* 2005; Martini *et al.* 2006; Wanat *et al.* 2008; Keelagher *et al.* 2011; Thacker *et al.* 2011). Class B mutants give NDD/RSD < 0.8 and include those with increased levels of precocious sister chromatid separation (PSCS) and/or meiosis II nondisjunction (MII-ND; e.g. *sgs1Δ*, *sgs1Δ795*, *cbf1Δ*; *iml3Δ*; and *cep1Δ*; Figure 4C; Masison and Baker 1992; Ghosh *et al.* 2004; Marston *et al.* 2004; Jessop *et al.* 2006; Rockmill *et al.* 2006; Chu and Burgess 2016). We found that the calculated NDD/RSD ratios for *nup2Δ*, *nup60Δ*, and *htz1Δ* were all less than 0.8 (Figure 4C; Table 5), which would classify them as Class B mutants.

To analyze the variation of the output values from TetFit-A, we ran 50 independent simulations using TetFit.Test using the calculated NDD and RSD values for each genotype and the corresponding number of experimentally dissected tetrads (Figure 4B, E columns; Table S2). The graphical output of the simulations shown in Figure 4C shows that each genotype gives well defined clusters representing NDD and RSD values for 50 simulated tetrad data sets.

## Discussion

Here we describe a systematic evaluation of growth and meiosis phenotypes caused by deleting nonessential Nups in the SK1 strain background. Comparisons for growth on rich media and minimal synthetic media at four-different temperatures uncovered growth phenotypes for *nup2Δ*, *nup60Δ*, *nup84Δ*, and *nup100Δ*. A growth defect for *nup84Δ* has been reported previously (Siniossoglou *et al.* 1996). For *nup60Δ* however, we found growth phenotypes (both positive and negative) where previous studies found no effect on growth on solid YPD media (Dilworth *et al.* 2005; Baryshnikova *et al.* 2010). Side-by-side comparisons would be necessary to conclude that the defect we see is due to the SK1 strain background.

The *nup2Δ*, *nup60Δ* and *nup100Δ* mutants showed increased fitness compared to WT under certain stress conditions. The *nup2Δ* and *nup60Δ* mutants formed larger colonies on YPD at 37° compared to WT, while *nup100Δ* mutant growth surpassed WT on minimal media at 25° and 30°. Increased fitness of a mutant compared to WT is characteristic of antagonistic pleiotropy (AP), where under certain environmental conditions, the WT copy of a gene decreases fitness compared to a mutant (Williams 2001). Genome-wide analysis of single-gene knockouts in yeast revealed that ~ 5.1% of nonessential genes show differences in the fitness of growth under one or more different challenged growth regimens (Qian *et al.* 2012). To our knowledge, the conditions where we found AP (i.e. 37° YPD and minimal media) have not been evaluated in genome-wide analysis (Baryshnikova *et al.* 2010; Qian *et al.* 2012). Our results indicate that Nup2, Nup60 and Nup100 suppress the efficiency of cell growth under certain conditions in the SK1 strain background.

Our analysis of sporulation and spore viability in the *nup* mutants uncovered phenotypes consistent with roles for Nup2, Nup60, Htz1 and Nup84 in the yeast meiotic program, but not Nup53, Nup100 and Nup157. We expected that these outcomes would correlate with the results of a genome-wide analysis of the yeast deletion collection that used a visual assay to score the segregation of a GFP-tagged chromosome III in intact tetrads (Marston *et al.* 2004). All of the mutants we analyzed were interrogated in this screen except for *nup157Δ*. While we found that spore viability of *nup53Δ* and *nup100Δ* and the kinetics of meiotic progression were the same as WT, the Marston study reported elevated levels of 2:2 GFP+: GFP-spores in these mutants, where primarily 4:0 would be expected for WT. In principle, the observed missegregtion of chromosomes would have resulted in increased spore death, but this was not the case for our analysis.

For *nup84Δ*, yet we found that less than 1% of these cells sporulated, while no decrease in sporulation or increased nondisjunction was reported in the Marston study. Dramatic differences like this could be due to the presence of a suppressor mutation that could have been introduced from the yeast deletion collection in the Marston study, for which there is precedence (Winzeler *et al.* 1999; Hughes *et al.* 2000). A more in depth evaluation of this phenotype could potentially uncover a functional link between these processes. For example, the meiotic defect we see in *nup84Δ* may reflect the defects of the *nup84Δ* mutant in telomere localization at the NE and DSB repair in vegetatively dividing cells as reported previously (Loeillet *et al.* 2005; Bukata *et al.* 2013).

For *nup6OΔ* and *htzlΔ* we found defects in spore viability. While spore inviability was not measured in the Marston study, they found no evidence for chromosome missegregation.

This would be consistent with the increased RSD we found for these mutants, however, TetFit estimated a modest contribution of MI-ND to spore death. On the other hand, Marston et al found that *nup2Δ* gave increased levels of 3:1 chromosome segregation errors, which is characteristic of defects leading to precocious sister chromatid separation (PSCS) or MII-ND segregation errors. This phenotype would appear as RSD using TetFit analysis and could possibly explain the increased spore death phenotype we saw for *nup2Δ*. Together, these results suggest that spore death in *htz1Δ* and *nup60Δ* is due to different causes than for *nup2Δ*.

The most surprising outcome of the study was that the *nup2Δ ndj1Δ* double mutant uniquely gave a synthetic block to completing the MI nuclear division compared to all of the other *nup ndj1Δ* double mutants. The *nup2Δ* mutation alone delayed progression through MI unlike *nup53Δ*, *nup100Δ,* and *nup157Δ*. These results suggest functional link between Nup2 and the chromosome events of meiosis. A more detailed analysis of Nup2's role in meiosis will be described in a future publication (D.B.C and S.M.B., unpublished).

## Materials and Methods

### Strains

All strains in this study are derivatives of SK1 and listed in Table 3. All media were generated as previously described (Treco and Lundblad 2001; Ho and Burgess 2011; Lui *et al.* 2013). All experiments were performed with at least two independent trials and in all cases similar results were obtained. Gene knockouts were constructed using standard tailed PCR based gene replacement and tagging techniques (Longtine *et al.* 1998; Goldstein and McCusker 1999).

**Table 3.**
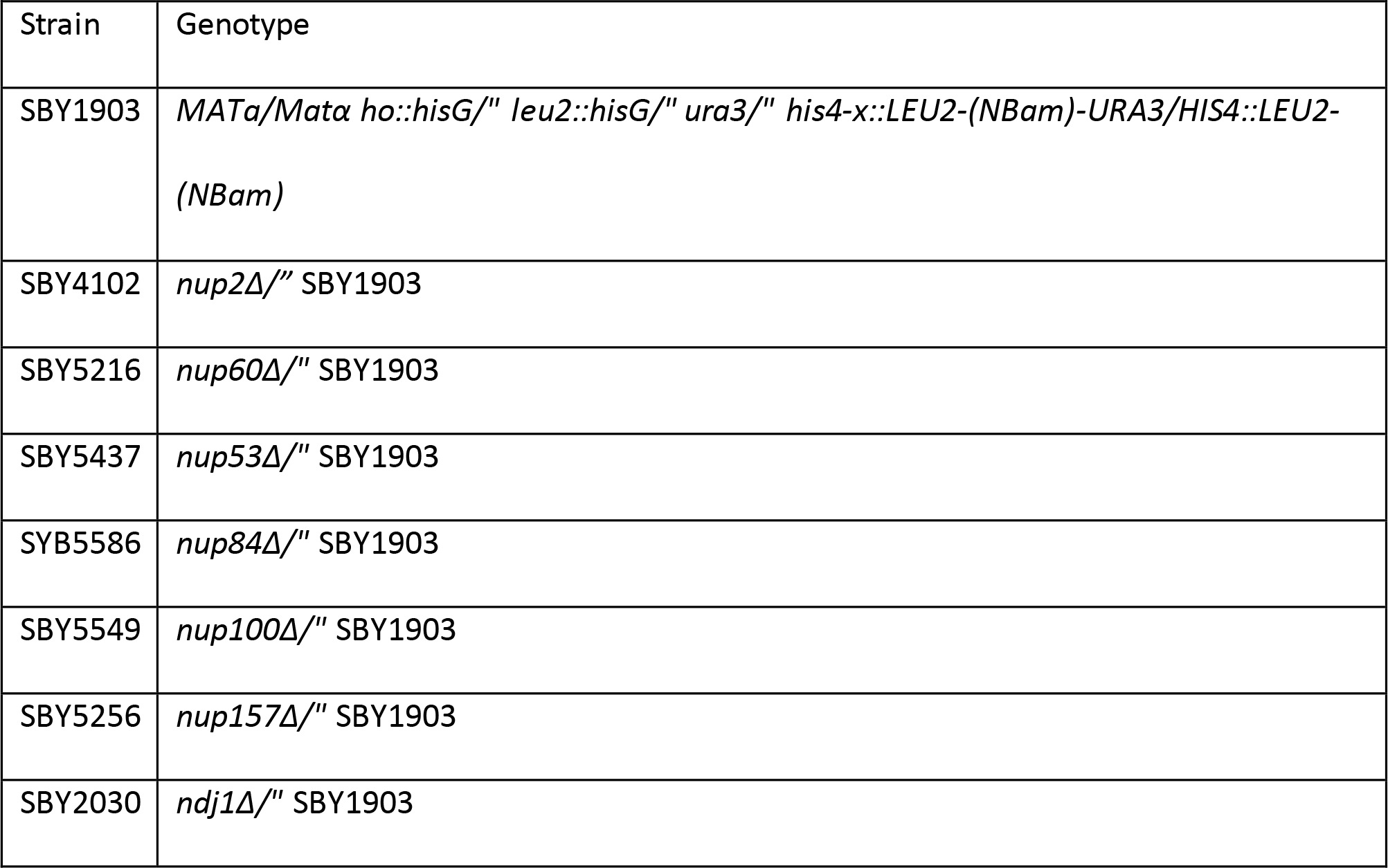
Yeast strains used in this study

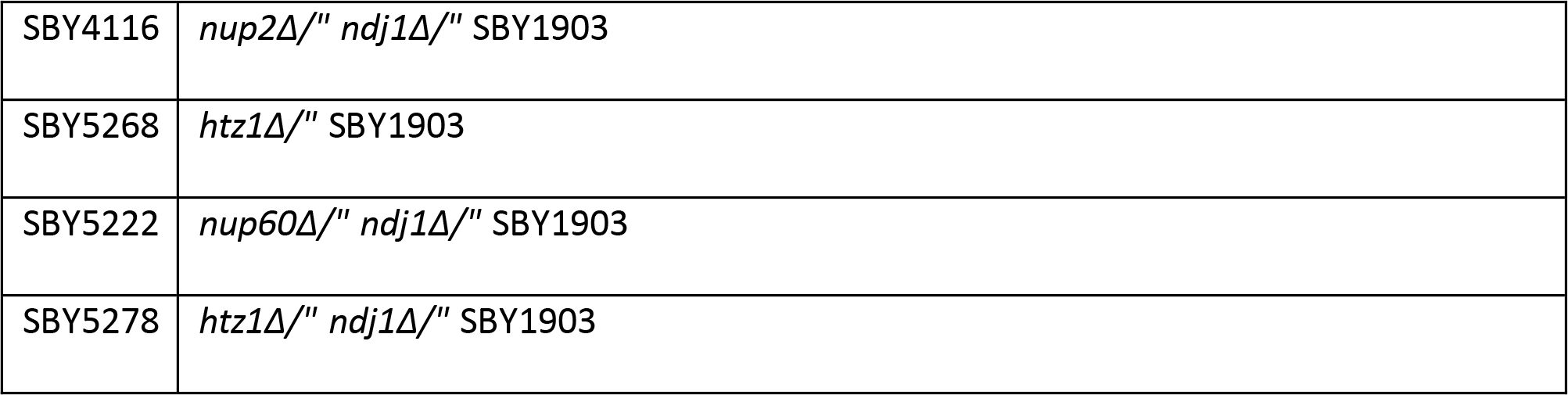

### Statistical Analysis

All statistical tests for significance are indicated in the text. Whenever multiple comparisons were performed the p values were corrected using the Holm method (Holm 1979).

### Spotting assay for vegetative growth

Diploid cells were prepared by first patching and mating haploid cells taken from glycerol stocks stored at - 80° on YPG plates (2% peptone, 1% yeast extract, 3% glycerol, 2% bacto agar, 0.01% adenine sulphate, 0.004% tryptophan, 0.002% uracil) for 15 hours at 30°. Cells were then streaked onto YPD plates (2% peptone, 1% yeast extract, 2% glucose, 2% bacto agar, 0.01% adenine sulphate, 0.004% tryptophan, 0.002% uracil) for 2 days at 30°. Well-isolated diploid single colonies were inoculated into 5ml liquid YPD (2% peptone, 1% yeast extract, 2% glucose, 0.01% adenine sulphate, 0.004% tryptophan, 0.002% uracil) for 24 hours at 30° on a roller drum. The cells were diluted from their 5 ml culture to 0D600 = 1.0. Suspended cells were serially diluted 1:10 in a 96-well plate and 2.5 |al of the diluted cultures were spotted onto YPD and minimal medium plates (1% Difco yeast nitrogen base without amino acids, 2% glucose, 2% bacto agar) using a multichannel pipettor, followed by incubation at 20°, 25°, 30°, and 37°. YPD plates were imaged after incubation for 48 and 72 hours and minimal medium plates were imaged after incubation for 72 and 120 hours using an Epson Perfection 1200 U scanner.

### Time course protocol

The time course protocol was followed as previously reported (Ho and Burgess 2011; Lui *et al.* 2013). Cells were synchronized for progression through meiosis by the same regimen used for assaying vegetative growth (above), except that the inoculated YPD liquid culture was grown for 30 hours instead of 24 hours and was not supplemented with uracil. Following the 30-hour growth period, cells were added to YPA (2% peptone, 1% yeast extract, 1% KOAc, 0.01% adenine sulphate, 0.004% tryptophan) to a final 0D600 of 0.23 and grown for 14.5 hours at 30° on a roller drum. Cells were pelleted at 1000 x *g* for 3 minutes, washed with SPM (1% KOAc, 0.02% raffinose, 0.009% synthetic complete dropout powder) and resuspended in SPM to a final 0D600 of 3.0. Cells were removed from the culture at various time points thereafter (starting at t = 0 hours), fixed in 40% ethanol and stained with DAPI to follow the kinetics of MI progression. The MI division was marked by the formation of cells with two well-differentiated DAPI stained foci. At least 200 cells were analyzed for each time point.

### Spore viability and nondisjunction analysis

Viability was determined as the percent viable spores from dissected tetrads. Single colonies from YPD plates were patched onto solid SPM media (1% KOAc, 0.02% raffinose, 0.009% synthetic complete dropout powder, 2% bacto agar), incubated for three days at 30°, and then dissected on YPD plates. The YPD plates were incubated for four days at 30° and colony formation was evaluated by unaided visual inspection. Any visible growth was considered to be a viable spore. The calculated MI-ND frequencies were determined using TetFit (Chu and Burgess 2016). Expected live:dead tetrad frequencies were generated for all genotypes using TetFit with the following conditions: the number of nondisjunction intervals (ndint) was 500, the number of random spore death intervals (rsdint) was 500, aneuploidy induced death (anid) was 0.035, and the MI-ND multiplier (ndm) was 10. Detailed methods are described in File S1.

### Data Availability

All strains are available upon request. File S1 contains a detailed description of the R-scripts used in this study and example data input and output formats associated with TetSim, TetFit and TetFit.Test.

## Acknowledgements

We thank JoAnne Engebrecht, Anne Britt, James McGehee, and An Nguyen for their insightful review of this manuscript. The authors have declared that no conflicting interests exist. This work was funded by NIH R01 GM075119 awarded to S.M.B.

## Abbreviations List

WT - Wild type

MI - Meiosis I

MII - Meiosis II

ND - Nondisjunction

NPC - Nuclear pore complex

Nup - NucleoporinReferences

## References

Aitchison,J. D., and M. P. Rout, 2012 The yeast nuclear pore complex and transport through it. Genetics 190: 855–883.

Aitchison,J. D., M. P. Rout, M. Marelli, G. Blobel and R. W. Wozniak, 1995 Two novel related yeast nucleoporins Nup170p and Nup157p: complementation with the vertebrate homologue Nup155p and functional interactions with the yeast nuclear pore-membrane protein Pom152p. J Cell Biol 131: 1133–1148.

Akiyama,K., J. Noguchi, M. Hirose, S. Kajita, K. Katayama et al., 2013 A mutation in the nuclear pore complex gene Tmem48 causes gametogenesis defects in skeletal fusions with sterility (sks) mice. J Biol Chem 288: 31830–31841.

Alber,F., S. Dokudovskaya, L. M. Veenhoff, W. Zhang, J. Kipper et al., 2007 The molecular architecture of the nuclear pore complex. Nature 450: 695–701.

Baryshnikova,A., M. Costanzo, Y. Kim, H. Ding, J. Koh et al., 2010 Quantitative analysis of fitness and genetic interactions in yeast on a genome scale. Nature methods 7: 1017–1024.

Bukata,L., S. L. Parker and M. A. D'Angelo, 2013 Nuclear pore complexes in the maintenance of genome integrity. Current opinion in cell biology 25: 378–386.

Cheslock,P. S., B. J. Kemp, R. M. Boumil and D. S. Dawson, 2005 The roles of MAD1, MAD2 and MAD3 in meiotic progression and the segregation of nonexchange chromosomes. Nat Genet 37: 756–760.

Chu,D. B., and S. M. Burgess, 2016 A computational approach to estimating nondisjunction frequency in Saccharomyces cerevisiae. G3: Genes | Genomes | Genetics: g3. 115.024380.

Chua,P. R., and G. S. Roeder, 1997 Tam1, a telomere-associated meiotic protein, functions in chromosome synapsis and crossover interference. Genes Dev 11: 1786–1800.

Church,K., 1976 Arrangement of chromosome ends and axial core formation during early meiotic prophase in the male grasshopper Brachystola magna by 3D, E.M. reconstruction. Chromosoma 58: 365–376.

Conrad,M. N., A. M. Dominguez and M. E. Dresser, 1997 Ndj1p, a meiotic telomere protein required for normal chromosome synapsis and segregation in yeast. Science 276: 1252–1255.

Denning,D., B. Mykytka, N. P. Allen, L. Huang, B. Al *et al.*, 2001 The nucleoporin Nup60p functions as a Gsp1p-GTP-sensitive tether for Nup2p at the nuclear pore complex. J Cell Biol 154: 937–950.

Dilworth,D. J., A. J. Tackett, R. S. Rogers, E. C. Yi, R. H. Christmas et al., 2005 The mobile nucleoporin Nup2p and chromatin-bound Prp20p function in endogenous NPC-mediated transcriptional control. J Cell Biol 171: 955–965.

Elrod,S. L., S. M. Chen, K. Schwartz and E. O. Shuster, 2009 Optimizing sporulation conditions for different Saccharomyces cerevisiae strain backgrounds. Methods Mol Biol 557: 21–26.

Fahrenkrog,B., W. Hubner, A. Mandinova, N. Pante, W. Keller et al., 2000 The yeast nucleoporin Nup53p specifically interacts with Nic96p and is directly involved in nuclear protein import. Mol Biol Cell 11: 3885–3896.

Fan,F., C. P. Liu, O. Korobova, C. Heyting, H. H. Offenberg et al., 1997 cDNA cloning and characterization of Npap60: a novel rat nuclear pore-associated protein with an unusual subcellular localization during male germ cell differentiation. Genomics 40: 444–453.

Ghosh,S. K., S.Sau, S. Lahiri, A. Lohia and P. Sinha, 2004 The Iml3 protein of the budding yeast is required for the prevention of precocious sister chromatid separation in meiosis I and for sister chromatid disjunction in meiosis II. Curr Genet 46: 82–91.

Gigliotti,S., G. Callaini, S. Andone, M. G. Riparbelli, R. Pernas-Alonso et al., 1998 Nup154, a new Drosophila gene essential for male and female gametogenesis is related to the nup155 vertebrate nucleoporin gene. J Cell Biol 142: 1195–1207.

Goldstein,A. L., and J. H. McCusker, 1999 Three new dominant drug resistance cassettes for gene disruption in Saccharomyces cerevisiae. Yeast 15: 1541–1553.

Ho,H. C., and S. M. Burgess, 2011 Pch2 acts through Xrs2 and Tel1/ATM to modulate interhomolog bias and checkpoint function during meiosis. PLoS Genet 7: e1002351.

Holm,P., 1977 Three-dimensional reconstruction of chromosome pairing during the zygotene stage of meiosis in Lilium longiflorum (thunb.). Carlsberg Res. Commun. 42: 103–151.

Holm,S., 1979 A simple sequentially rejective multiple test procedure. Scandinavian journal of statistics: 65–70.

Hughes,T. R., C. J. Roberts, H. Dai, A. R. Jones, M. R. Meyer et al., 2000 Widespread aneuploidy revealed by DNA microarray expression profiling. Nat Genet 25: 333–337.

Jessop,L., B. Rockmill, G. S. Roeder and M. Lichten, 2006 Meiotic chromosome synapsis-promoting proteins antagonize the anti-crossover activity of sgs1. PLoS Genet 2: e155.

Kane,S. M., and R. Roth, 1974 Carbohydrate metabolism during ascospore development in yeast. J Bacteriol 118: 8–14.

Keelagher,R. E., V. E. Cotton, A. S. Goldman and R. H. Borts, 2011 Separable roles for Exonuclease I in meiotic DNA double-strand break repair. DNA Repair (Amst) 10: 126–137.

Kellis,M., B. W. Birren and E. S. Lander, 2004 Proof and evolutionary analysis of ancient genome duplication in the yeast Saccharomyces cerevisiae. Nature 428: 617–624.

Kirkpatrick,D. T., J. R. Ferguson, T. D. Petes and L. S. Symington, 2000 Decreased meiotic intergenic recombination and increased meiosis I nondisjunction in exo1 mutants of Saccharomyces cerevisiae. Genetics 156: 1549–1557.

Klein,F., T. Laroche, M. E. Cardenas, J. F.-X.Hofmann, D. Schweizer et al., 1992 Localization of RAP1 and topoisomerase II in nuclei and meiotic chromosomes of yeast. The Journal of Cell Biology 117: 935–948.

Loeb,J., L. Davis and G. Fink, 1993 NUP2, a novel yeast nucleoporin, has functional overlap with other proteins of the nuclear pore complex. Molecular biology of the cell 4: 209–222.

Loeillet,S., B. Palancade, M. Cartron, A. Thierry, G.-F. Richard et al., 2005 Genetic network interactions among replication, repair and nuclear pore deficiencies in yeast. DNA repair 4: 459–468.

Longtine,M. S., A. McKenzie, 3rd, D. J. Demarini, N. G. Shah, A. Wach et al., 1998 Additional modules for versatile and economical PCR-based gene deletion and modification in Saccharomyces cerevisiae. Yeast 14: 953–961.

Lui,D. Y., C. K. Cahoon and S. M. Burgess, 2013 Multiple opposing constraints govern chromosome interactions during meiosis. PLoS Genet 9: e1003197.

Marelli,M., J. D. Aitchison and R. W. Wozniak, 1998 Specific binding of the karyopherin Kap121p to a subunit of the nuclear pore complex containing Nup53p, Nup59p, and Nup170p. The Journal of cell biology 143: 1813–1830.

Marston,A. L., W. H. Tham, H. Shah and A. Amon, 2004 A genome-wide screen identifies genes required for centromeric cohesion. Science 303: 1367–1370.

Martini,E., R. L. Diaz, N. Hunter and S. Keeney, 2006 Crossover homeostasis in yeast meiosis. Cell 126: 285–295.

Masison,D. C., and R. E. Baker, 1992 Meiosis in Saccharomyces cerevisiae mutants lacking the centromere-binding protein CP1. Genetics 131: 43–53.

Ptak,C., J. D. Aitchison and R. W. Wozniak, 2014 The multifunctional nuclear pore complex: a platform for controlling gene expression. Curr Opin Cell Biol 28: 46–53.

Qian,W., D. Ma, C. Xiao, Z. Wang and J. Zhang, 2012 The genomic landscape and evolutionary resolution of antagonistic pleiotropy in yeast. Cell reports 2: 1399–1410.

Rockmill,B., K. Voelkel-Meiman and G. S. Roeder, 2006 Centromere-proximal crossovers are associated with precocious separation of sister chromatids during meiosis in Saccharomyces cerevisiae. Genetics 174: 1745–1754.

Rout,M. P., J. D. Aitchison, A. Suprapto, K. Hjertaas, Y. Zhao et al., 2000 The yeast nuclear pore complex composition, architecture, and transport mechanism. The Journal of cell biology 148: 635–652.

Shonn,M. A., R. McCarroll and A. W. Murray, 2000 Requirement of the spindle checkpoint for proper chromosome segregation in budding yeast meiosis. Science 289: 300–303.

Siniossoglou,S., C. Wimmer, M. Rieger, V. Doye, H. Tekotte et al., 1996 A novel complex of nucleoporins, which includes Sec13p and a Sec13p homolog, is essential for normal nuclear pores. Cell 84: 265–275.

Smitherman,M., K. Lee, J. Swanger, R. Kapur and B. E. Clurman, 2000 Characterization and targeted disruption of murine Nup50, a p27(Kip1)-interacting component of the nuclear pore complex. Mol Cell Biol 20: 5631–5642.

Thacker,D., I. Lam, M. Knop and S. Keeney, 2011 Exploiting spore-autonomous fluorescent protein expression to quantify meiotic chromosome behaviors in Saccharomyces cerevisiae. Genetics 189: 423–439.

Treco,D., and V. Lundblad, 2001 Preparation of yeast media. Current protocols in molecular biology/edited by Frederick M. Ausubel…[et al.]: Unit13. 11.

Trelles-Sticken,E., J. Loidl and H. Scherthan, 1999 Bouquet formation in budding yeast: initiation of recombination is not required for meiotic telomere clustering. J Cell Sci 112 (Pt 5): 651–658.

Wanat,J. J., K. P. Kim, R. Koszul, S. Zanders, B. Weiner et al., 2008 Csm4, in collaboration with Ndj1, mediates telomere-led chromosome dynamics and recombination during yeast meiosis. PLoS Genet 4: e1000188.

Wente,S. R., M. P. Rout and G. Blobel, 1992 A new family of yeast nuclear pore complex proteins. The Journal of Cell Biology 119: 705–723.

Williams,G. C., 2001 Pleiotropy, natural selection, and the evolution of senescence. Science's SAGE KE 2001: 13.

Winzeler,E. A., D. D. Shoemaker, A. Astromoff, H. Liang, K. Anderson et al., 1999 Functional characterization of the S. cerevisiae genome by gene deletion and parallel analysis. Science 285: 901–906.

Zickler,D., and N. Kleckner, 1998 The leptotene-zygotene transition of meiosis. Annu Rev Genet 32: 619–697.

Zickler,D., and N. Kleckner, 2015 Recombination, Pairing, and Synapsis of Homologs during Meiosis. Cold Spring Harb Perspect Biol 7.

